# Actors’ Facial Movement Magnitude and Cardiac Dynamics Predict Observers’ Emotion Believability Ratings

**DOI:** 10.64898/2026.07.01.735852

**Authors:** Alejandro Galvez-Pol, Victoria Rambaud, Julia F. Christensen, James M. Kilner

## Abstract

In non-verbal communication, observers infer emotions from visible facial movements, yet emotional experiences are described in internal bodily terms (e.g., “my heart skipped a beat”)^1–3^. This contrast highlights a tension between external sensory cues and internal signals. In this context, we examined an overlooked gap in affective science: what makes an emotional portrayal believable, and do believability judgments reflect only what observers can see or also the portraying person’s internal cardiac dynamics? To test this, we created 311 scenario-driven acting clips designed to avoid prototypical posed displays^4–6^. For each clip, we quantified facial movement magnitude from the video, recorded ECG during preparation and enactment, and collected actors’ self-reports. Online participants (N = 371) viewed these clips and provided emotion recognition responses and continuous ratings of believability, valence, or arousal. The results show that believability decreased as movement magnitude increased, with a non-linear relationship indicating a stronger penalty as motion increased. Valence further shaped this pattern, with increasing movement reducing believability more strongly for portrayals with negative valence. This effect persisted after accounting for intended emotion, perceived arousal, and emotion recognizability. Cardiac dynamics varied during performance, and actors’ higher heart rate variability was associated with higher believability for positively valenced portrayals. Together, these findings show that believability is driven by visible movement cues interpreted in relation to valence, with actors’ cardiac dynamics showing selective alignment with believability. These results identify core components of believable emotional expressions and provide a basis for studying such judgments in everyday social interaction.

**Highlights:**

- Larger facial movements were associated with lower believability, non-linearly
- Negative valence portrayals were penalized more strongly for large movements
- Emotion recognizability did not explain believability, even after adjustment
- Actor heart-rate variability was associated with higher believability in positive portrayals

## Results

Actors performed short scenario scripts and enacted target emotions in a self-paced setting. Because no available stimulus set combined acted portrayals with movement, physiology, self-report, and observer data, we created a dedicated stimulus set (Figure 1; STAR Methods)^6–9^. For each clip, we extracted a stimulus-level movement magnitude measure from the video and recorded electrocardiogram (ECG) during preparation and enactment of the portrayals. Online participants provided emotion recognition responses and ratings of believability, valence, and arousal. The resulting clips occupied distinct but partially overlapping regions of affective space defined by independent valence and arousal ratings (Figure S1). We used these measures to test whether facial movement magnitude was associated with believability at the stimulus and trial level, whether valence modulated this relationship, whether recognizability accounted for believability, and whether cardiac dynamics during performance showed any correspondence with observer judgments.

**Figure 1.**
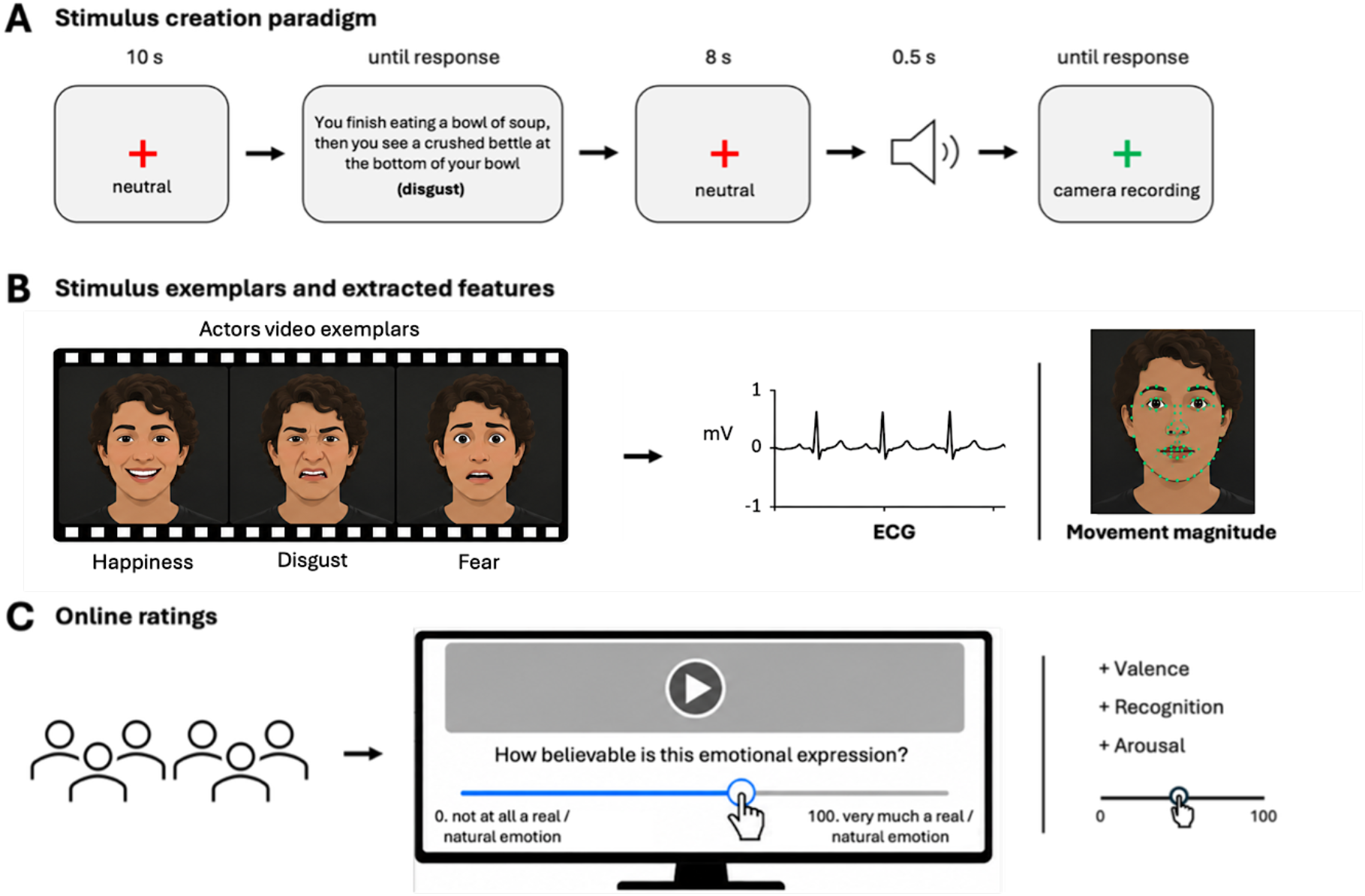
Stimulus creation, feature extraction, and online validation. **(A) Stimulus creation paradigm**. Each trial began with a fixation cross on the screen, followed by a short scenario describing an emotional instance (example shown). Actors then enacted the target emotion while facing the camera. Enactments were self-paced and recorded as dynamic facial-expression video clips. **(B) Stimulus outputs and extracted measures**. The resulting stimuli set comprised multiple actors and emotion categories (filmstrip illustrates exemplars). We extracted ECG-derived cardiac measures during preparation and enactment of each portrayal (example ECG trace shown) and quantified facial movement magnitude from the video using the OpenFace tool, yielding stimulus-level behavioral and physiological descriptors. **(C) Stimulus online validation and ratings**. Prolific-based participants rated the stimuli in an online experiment. Each participant viewed a batch of clips and provided continuous ratings of believability, valence, and arousal, and completed an emotion-recognition task (forced choice, including an “other” option). Ratings of believability, valence, or arousal were collected via slider responses. ***Preprint note:*** *Faces shown in (B) are synthetic and do not depict the actual study actors. This substitution was made in compliance with bioRxiv’s policy on identifiable images*.

### Increased facial movements reduce perceived believability

We first quantified actors’ facial motion as movement magnitude and related it, at the stimulus level, to mean observer believability (0–100) across 304 clips. The results of this analysis showed that perceived believability decreased as movement magnitude increased (r = –0.463, 95% CI [–0.547, –0.370], p < 0.001). To express the effect size on the believability scale, we next regressed mean believability on z-scored movement magnitude (coefficients reflecting a 1 SD change). This analysis showed that a 1 SD increase in movement magnitude predicted a decrease of 5.8 in mean believability (β = –5.835, R^2^ = 0.214, p < 0.001). Moreover, the association persisted after controlling for the intended emotion category that actors portrayed (β = –6.211, R^2^ = 0.443, p < 0.001), indicating that it was not merely driven by differences between emotion categories. Trial-level mixed-effects analyses reported next provide the primary inferential test while accounting for repeated ratings within observers and clips.

We then asked whether this movement-to-believability association was strictly linear. Here a quadratic model fit better than a linear model, with a negative linear term (β = –9.586, p < 0.001) and a positive quadratic term (β = 1.456, p < 0.001; R^2^ = 0.252). This indicates that as movement magnitude increased, portrayals were increasingly judged as less believable, with the penalty strongest as movement increased beyond the range associated with higher believability^5,10–13^. Larger facial motion is therefore associated with less believable portrayals, and the relationship is not fully linear.

### Movement relates to believability beyond affective context

We next tested whether the previously described movement effect generalized at the trial level. Because facial movements are interpreted in relation to affective context, including intended emotion, arousal, and valence, we asked whether movement magnitude remained associated with believability after accounting for these factors^14–16^. Using mixed-effects models with random intercepts for observer and stimulus, we accounted for the fact that each clip was rated multiple times, and each observer rated multiple clips (11,323 ratings, 93 believability raters, 304 clips). Here, incorporating intended emotion (the target category the actor aimed to portray) improved the model beyond movement magnitude alone (χ^2^(6) = 104.66, p < 0.001), and adding stimulus-level mean arousal and mean valence further improved model fit (χ^2^(2) = 113.09, p < 0.001).

Crucially, movement magnitude remained robustly associated with believability in the full model. Higher movement magnitude predicted lower believability (β = –4.462, SE = 0.611, p < 0.001). Mean arousal also predicted lower believability (β = –3.757, SE = 0.830, p < 0.001), whereas mean valence predicted higher believability (β = 10.288, SE = 1.067, p < 0.001). Hence, even after accounting for which emotion was intended and how arousing or positive the clips were perceived to be, larger facial movements were still associated with lower believability.

### Valence modulates the non-linear movement-to-believability relationship

Because facial expressions convey both emotion categories and affective dimensions, the valence of a portrayal may shape how observers evaluate whether visible facial dynamics are believable^16,17^. Given the non-linear association observed at the stimulus level, we tested whether a more flexible model captured the trial-level movement–believability relationship and whether this relationship varied with perceived valence^17,18^. Here, a non-linear spline model described the trial-level data better than a linear model (χ^2^(2) = 10.952, p = 0.004).

We then allowed the movement-to-believability function to vary with stimulus-level valence. The results of this analysis explained additional variation (χ^2^(3) = 15.211, p = 0.002), indicating that the same movement magnitude can be judged differently depending on valence. In more negatively valenced portrayals, increased movement was penalized more strongly, whereas in more positively valenced portrayals comparable movement remained more believable. Figure 2 visualizes this relationship and how it changes with valence.

**Figure 2.**
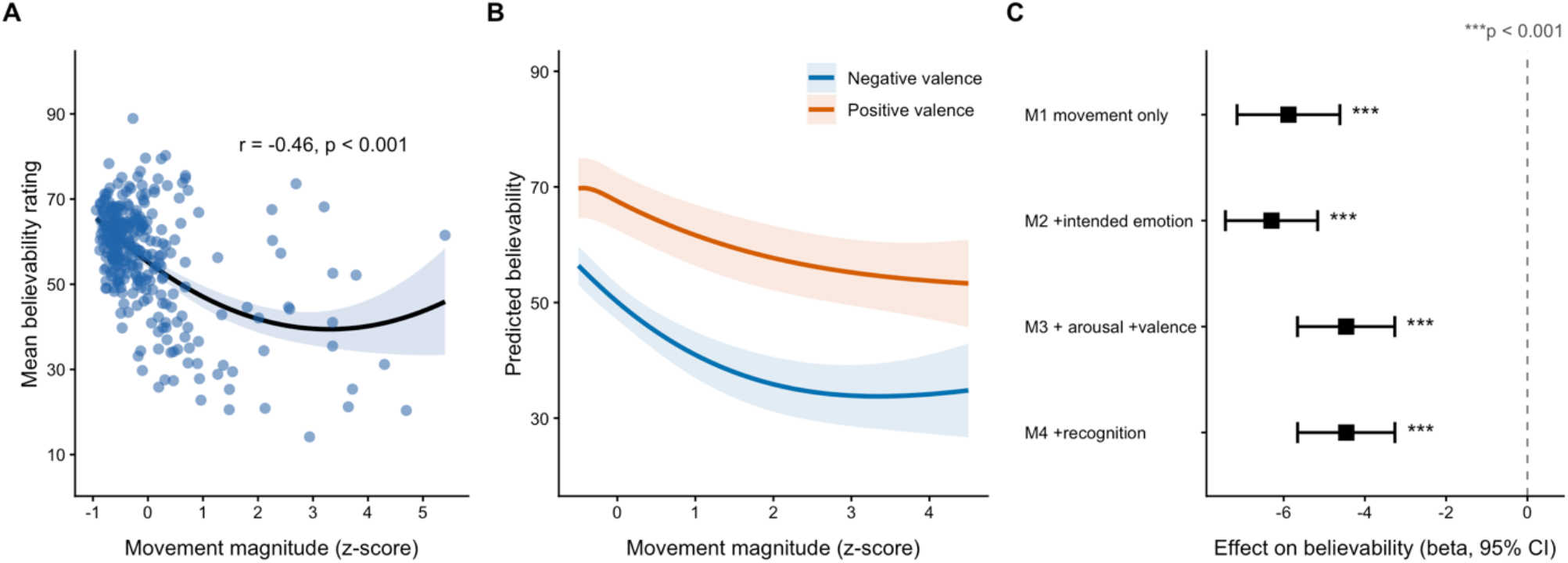
Facial movement magnitude predicts believability through a non-linear relationship modulated by valence. **(A)** Stimulus-level relationship between facial movement magnitude (z-scored across stimuli) and mean believability (0–100) across stimuli. The line shows the quadratic fit; the shaded band indicates the 95% CI. **(B)** Trial-level non-linear movement–believability relationship from a linear mixed-effects model with natural spline basis functions interacted with stimulus-level valence. Curves show predicted believability at valence_z = −1 and +1 (negative, positive portrayals); arousal held at its mean and predictions averaged across intended emotion categories; random intercepts for observer and stimulus. Shaded bands: 95% Wald CIs. **(C)** Stability of the movement effect across nested linear mixed-effects models of trial-level believability (random intercepts for observer and stimulus). Squares show the fixed-effect estimate for movement magnitude (β) with 95% CIs as covariates are added sequentially: intended emotion, mean arousal and mean valence, and recognizability.

Finally, we asked whether this context dependence reflected valence specifically, or whether emotion categories (operationalized as intended emotion) added further explanatory value. Discrete intended emotion provided additional modulation beyond valence (χ^2^(18) = 34.355, p = 0.011; Figure S2), although the improvement in fit was modest relative to the valence effect, suggesting that affective dimensions rather than discrete categories primarily drive the context dependence of the movement–believability mapping.

### Recognition accuracy does not explain believability

Many widely used facial-expression stimulus sets were tested for recognition accuracy, although recognizability and perceived believability are not equivalent dimensions of expression perception^6,19–22^. We therefore tested whether emotion recognizability related to perceived believability. At the stimulus level, emotion recognizability (i.e., proportion of participants choosing the actor’s intended emotion) showed a modest negative association with mean believability (r = −0.186, 95% CI [−0.292, −0.075], p = 0.001). However, this association was not evident after adjusting for movement magnitude, mean arousal, and mean valence (β = −0.182, p = 0.705). Consistently, adding recognizability as a stimulus-level covariate to the trial-level mixed model did not improve model fit (χ^2^(1) = 0.210, p = 0.647), and the movement magnitude effect remained virtually unchanged (β = −4.462 vs. −4.460). Thus, a portrayal can be readily identifiable in terms of which emotion it conveys without necessarily appearing believable.

### Actors’ cardiac dynamics show valence-dependent alignment with observer believability

To assess whether observer believability judgments related to actor-centered measures beyond their visible movement, we analyzed actors’ cardiac dynamics (heart rate and HRV). Because emotional episodes are accompanied by changes in peripheral physiology, and HRV has been linked to social-affective processing^2,23–28^, we first tested whether heart rate differed across intended emotions during the preparation and enactment periods of the portrayals. Here we computed heart rate for each stimulus relative to the actor’s own mean (actor-mean corrected heart rate). Heart rate during preparation of the portrayals did not differ between intended emotions (F(6, 281) = 0.94, p = 0.464), whereas heart rate during the enactment of portrayals differed significantly between intended emotions (F(6, 288) = 12.662, p < 0.001; Figure 3A). Consistent with this heart-rate enactment effect, the preparation-to-enactment change in heart rate also varied by intended emotions (F(6, 281) = 6.238, p < 0.001), indicating systematic emotion-dependent cardiac modulation during enactment.

**Figure 3.**
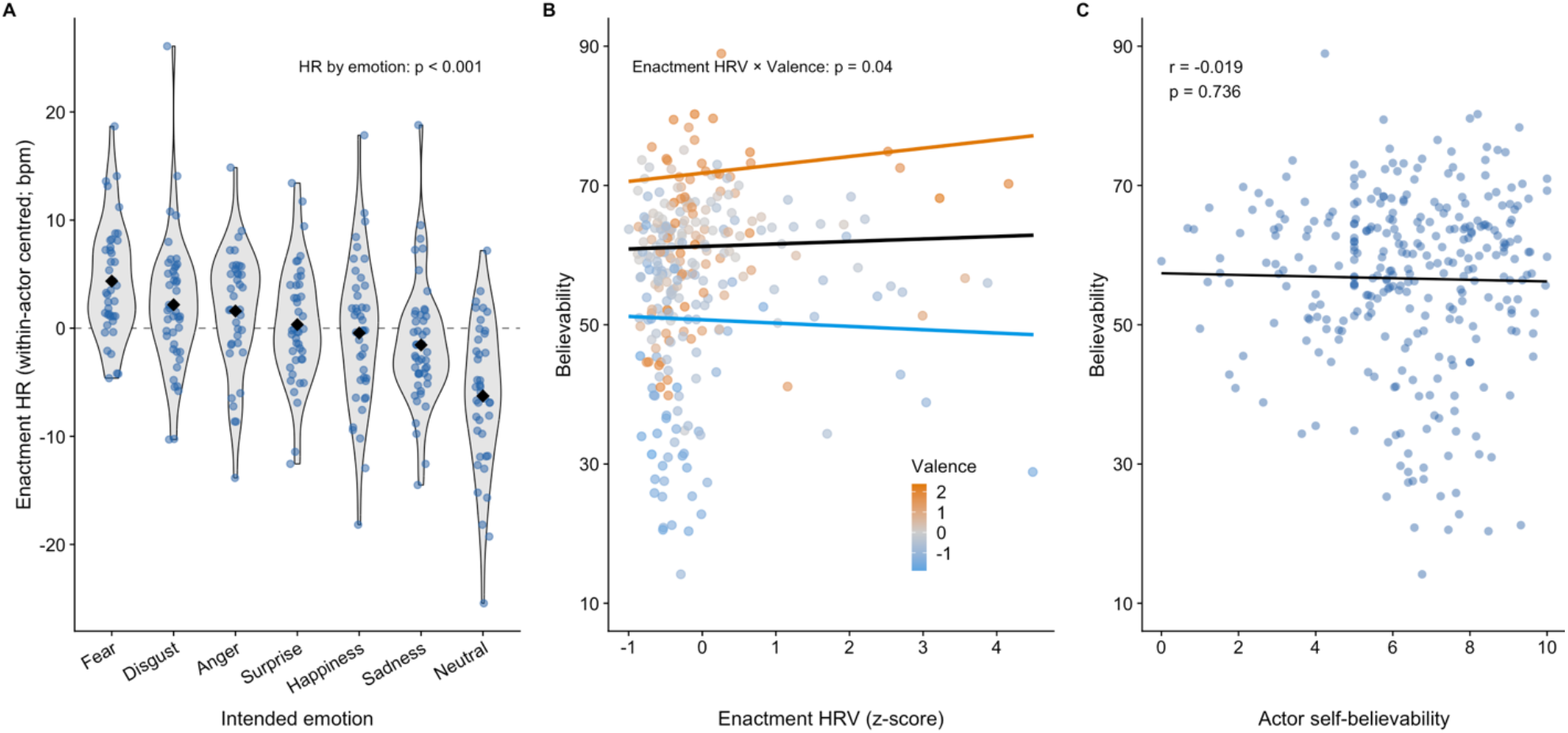
Cardiac dynamics vary during acting and align selectively with believability. **(A)** Enactment heart rate by intended emotion, expressed relative to each actor’s average heart rate (HR) across stimuli. Each point is a stimulus; violins show the distribution; the black diamond indicates the mean per emotion. Enactment HR differed across intended emotions (p < 0.001). **(B)** Enactment HRV (z-scored across stimuli) versus mean observer believability (y). Each point is a stimulus, colored by stimulus-level valence on a continuous scale from more negative to more positive. Lines show modelled HRV–believability relationships at mean valence (black) and at valence_z = −1 and +1 (blue and orange), based on the trial-level mixed model including movement magnitude, mean arousal, intended emotion, and an HRV × valence interaction. This pattern indicates a valence-dependent HRV–believability association, with higher HRV associated with higher believability under more positive valence. **(C)** Mean observer believability plotted against actor self-rated believability across stimuli. Actor self-believability did not correlate with observer believability (r = −0.019, p = 0.736), indicating limited actor– observer correspondence.

We next asked whether HRV during preparation and enactment was associated with observers’ believability judgments. Using the same set of 285 clips with complete preparation and enactment HRV data, preparation and enactment HRV each showed valence-dependent associations with observer believability ratings (HRV × valence interactions, all ps ≤ 0.040). To interpret these interactions, we estimated the HRV–believability slope at representative points along the continuous valence scale (Figure 3B). At one SD above the mean valence (valence_z = +1), higher HRV predicted higher believability for enactment HRV (p = 0.026) and as a trend for preparation HRV (p = 0.080). At one SD below the mean (valence_z = −1), no such association emerged for either measure (prep p = 0.076, enact p = 0.479). The association between HRV and believability became more positive as portrayals increased in valence. This pattern was clearest for enactment HRV, where higher HRV predicted higher believability at positive valence, and was not evident at negative valence. This pattern was not driven by facial movement magnitude as it persisted in models that additionally allowed movement magnitude to interact with valence (Figure S3), ruling out the possibility that the HRV effect merely reflected differences in facial movement magnitude.

Taken together, these results indicate that cardiac dynamics vary systematically during performance, but their association with observer believability is selective. Higher HRV was associated with higher believability for more positively valenced portrayals (*vs*. more negative), particularly for enactment HRV. This pattern extends empirical work linking a higher HRV to a better skill in conveying some facial expressions^29^ by showing that cardiac dynamics also relate to observers’ believability judgments, and that this relation depends on the emotional valence of the portrayal.

### Exploratory analysis: actors’ subjective reports diverge from observers’ experience

Finally, because believability-related judgments are typically treated as observer-side inferences rather than direct readouts of the actor’s own experience^30,31^, we tested whether actors’ self-assessments aligned with how believable their performances appeared to observers. Actor self-believability did not correlate with mean observer believability (r =−0.019, p = 0.736), and self-reports of emotional strength added no explanatory value to the believability model (χ^2^(1) = 0.010, p = 0.971). Mediation analyses revealed a specific indirect pathway. Actors who self-reported greater emotional strength tended to produce slightly less facial movement (path a: β = −0.094, p = 0.049), and clips with less facial movement were rated as more believable by observers (path b: β = −4.465, p < 0.001), yielding a positive indirect effect (indirect effect = 0.415, 95% CI [0.084, 0.804], B = 5000 bootstrap replicates). However, the total effect of actor-reported emotional strength on observer believability was non-significant (χ^2^(1) = 0.515, p = 0.473), indicating that this indirect pathway did not translate into any detectable influence on how believable the performance appeared to observers. Together, these results point to a gap between actors’ self-assessments and observer experience. Perceived believability appears to be driven by visible movement cues interpreted in relation to affective meaning and the actors’ internal cardiac dynamics, but not by how intensely the actor experienced the portrayed emotion.

## Discussion

Believability is a core constraint in professional acting as much as in general social communication, yet it has received comparatively little empirical attention relative to emotion recognition accuracy and emotion categorization^6,32,33^. Our results indicate that believability judgments follow plausibility criteria that depend strongly on the magnitude of visible facial movement and on valence. This distinction matters because an emotional portrayal can be correctly recognized while still being judged as unbelievable, exaggerated, staged, or strategically performed, shaping later inferences about emotion, intent, and trust^12,30,34^. Across portrayals in this experiment, larger movement magnitude was associated with lower believability, and this association was non-linear (quadratic, in fact), with the decline occurring once motion departed from naturalistic expectations. This emphasis on dynamic movement is consistent with recent work showing that facial expressions are processed as time-varying social signals, with dynamic movement contributing to emotion perception and its neural representation^8,13,35^.

Valence further calibrated this movement-believability relationship. Larger facial movement magnitude was penalized more strongly in negatively valenced portrayals than in positively valenced portrayals. Intended emotion also provided additional, although comparatively modest, modulation beyond valence, suggesting that both broad affective dimensions (i.e., valence, but not arousal) and specific intended emotion shape how movement is perceived. Actors’ cardiac dynamics (heart rate, HR, and heart rate variability, HRV) measured during performance showed a related but more selective pattern. Heart rate modulation from preparation to enactment varied with intended emotion, indicating systematic physiological adjustment during performance. HRV also aligned with believability, but primarily in positively valenced portrayals, and this association remained after accounting for facial movement magnitude. Together, these findings suggest that believability is not fully captured by visible facial movement or cardiac dynamics in isolation. Instead, both measures relate to believability in ways that depend on the affective context of the portrayal^3,17,36^.

A central motivation of the study was the gap between what observers see when an actor expresses an emotion and the internal bodily signals people often invoke when describing an emotion they are having (e.g., a racing heart). The findings from our cardiac dynamics analysis suggest that internal dynamics are engaged during actors’ emotional portrayals, in accordance with the bodily signals described in the above example. However, these cardiac dynamics do not yield a stable “believability signature” that observers can rely on across emotional portrayals. Instead, the physiology-believability association was selective: higher HRV was associated with greater believability mainly for positively valenced portrayals. This pattern extends empirical work linking HRV to deliberate facial expressive skill and complements evidence that cardiac signals can contribute to social-affective perception^25,29,37–41^. At the same time, it cautions against treating physiology as a general proxy for believability. Internal bodily dynamics may contribute to believable performance, but their relevance appears to depend on the affective frame (valence) in which the expression is produced and evaluated.

The movement and valence effects are consistent with a conservative plausibility gate for social signals^7,31,42,43^. Highly amplified movement may reduce believability when it exceeds observers’ expectations about how an emotion should plausibly unfold. The stronger penalty for larger movement magnitude in negatively valenced portrayals fits this account, because contexts associated with threat, conflict, or distress may place stronger constraints on what counts as a believable expression. Consistent with this observer-centered interpretation, actors’ self-believability and emotional strength showed little correspondence with observer believability. Thus, believability judgments appear to be shaped more by the visible expressive signal and valence than by the actor’s own evaluation of how believable they were. The physiology findings add a further constraint: if internal bodily dynamics were transparently expressed and reliably read out by observers, one would expect a more uniform physiological correspondence with believability. Instead, the HRV pattern suggests that any contribution of internal state information is partial, valence dependent, and likely constrained by the plausibility of visible expressive dynamics.

In terms of limitations, several boundaries constrain the generality of these findings^44^. First, because the stimuli were acted portrayals, the extent to which the same believability criteria generalize to spontaneous emotional behavior in everyday interaction remains to be tested^4,7,22^. Second, the stimulus set was produced by a limited number of actors and rated by online observers, so replication across a broader range of performers and populations would strengthen generality. Third, cardiac measures are indirect indices of internal state, and the observed HRV–believability alignment was selective, so stronger mechanistic conclusions about how physiology is expressed in performance would be premature.

A next step is to test whether the same movement-based believability criteria generalize to non-performance settings in which believability impressions are more implicit, and to refine the physiology component by examining additional autonomic features and temporal dynamics that may be informative. Beyond actors’ physiology, observers’ own cardiac dynamics during stimulus viewing may also shape believability judgments, as cardiac signals have been shown to facilitate the breakthrough of emotional stimuli into conscious awareness^45^, raising the possibility that an observer’s cardiac phase or state could modulate sensitivity to subtle believability cues. For instance, respiration is a consciously accessible autonomic signal that could shape both expressive timing and observers’ impressions of believability, making it a promising target for testing how bodily dynamics contribute to convincing emotional expression^40,46^.

## Conclusions

A smile, a grimace, and a flinch can each be easy to recognize, yet still feel unconvincing. Here we show that believability is shaped primarily by visible facial movement, specifically by how far movement departs from a plausible range, and that this relationship shifts with emotional valence. Actors’ cardiac dynamics varied during different emotional portrayals, and aligned with observer believability ratings specifically for positive portrayals, suggesting that actors’ internal bodily dynamics may relate selectively to observers’ believability ratings. More broadly, the findings tap into a fundamental challenge in emotion perception: observers judge visible facial movements, while many bodily changes associated with emotion remain hidden from view. Our results suggest that facial expressions are not direct readouts of a person’s internal state, but context-embedded social signals whose believability depends on visible behavior, affective meaning, and selective bodily correspondence. This positions believability as a measurable judgment in social perception, with relevance for understanding how observers evaluate emotional expressions across acted, spontaneous, and real-world settings.

## RESOURCE AVAILABILITY

### Lead contact

Further information and requests for resources should be directed to and will be fulfilled by the lead contact, Alejandro Galvez-Pol.

### Materials availability

The video stimulus set generated in this study will not be made publicly available because the clips contain identifiable actors performing emotional portrayals, and the accompanying ratings include believability judgments that could be misused to rank, compare, or evaluate individual performers. To protect the actors from potential reputational or professional harm, access to the video materials will be controlled by the lead contact. Researchers wishing to use the clips may request access from the lead contact and will be required to sign a materials-use agreement. This agreement will specify that the materials may be used only for approved research purposes; that individual actors must not be named, ranked, profiled, or described as more or less believable, authentic, skilled, or convincing; that results must be reported only at aggregate or anonymized levels; and that the clips, ratings, and actor-linked metadata must not be redistributed, used for commercial evaluation, or used in any context that could affect the performers’ reputation or career opportunities.

### Data and code availability

Analysis code and non-identifying behavioral data needed to reproduce the reported statistical analyses will be made available on OSF upon publication. An anonymized view-only link is available for peer review. Access to video stimuli and actor-linked metadata is restricted for ethical reasons described under Materials availability and can be provided only under a controlled materials-use agreement.

## ACKNOWLEDGMENTS

We thank all actors and actresses who participated in this study for their time, commitment, and generosity in contributing the emotional portrayals that made this work possible. We also thank all participants and colleagues who helped to advance the study. This research was supported by the Leverhulme Trust, United Kingdom (grant RPG-2016-120).

## AUTHOR CONTRIBUTIONS

**A.G.-P.: Conceptualization, Methodology, Validation, Formal analysis, Investigation, Data curation, Writing – original draft, Writing – review and editing, Visualization, Supervision, Project administration, Funding acquisition. V.R.: Software, Formal analysis, Data curation, Visualization, Writing – review and editing. J.F.C.: Conceptualization, Methodology, Validation, Investigation, Writing – review and editing. J.M.K.: Conceptualization, Methodology, Supervision, Writing – review and editing. All authors reviewed and approved the final manuscript**.

## Methods

### Ethics approval and consent

Data collection was approved by the UCL Research Ethics Committee / Queen Square Institute of Neurology ethics committee. All actors provided written informed consent to take part in the study and separately consented to the use of their images (including motion images) for research purposes and publication in scientific communications, with the explicit condition that their name would not be linked to the materials. Online rating participants provided informed consent prior to participation.

### Stimulus set overview

Three-hundred-eleven video clips of emotional facial expressions (duration 6–13 s; M = 10.9, SD = 2.1; 24 fps) were created for this investigation. Fifteen professional actors enacted seven basic emotions (happiness, sadness, surprise, disgust, anger, fear, neutral). Portrayals were dynamic, scenario-driven, with the actors looking directly into the camera, and expressing emotions non-verbally.

### Scenario scripts

Emotional portrayals were elicited by scripted scenarios. We initially generated a pool of 70 emotion-eliciting scripts, each consisting of a neutral context sentence followed by a second sentence describing an event linked to a target emotion. To validate script–emotion correspondence, 27 volunteers completed an online Qualtrics® survey and selected which emotion (six basic emotions plus “other”) best matched each scenario. We then created 10 additional neutral scenarios and recruited a second group of 16 volunteers to label the emotion associated with the selected scenarios and the neutral scenarios (six basic emotions, neutral, or “other”). For recording, we selected the three most frequently endorsed scenarios for each of the six emotions and for neutral, yielding 21 scenarios in total (18 emotional, 3 neutral).

### Actor recruitment and preparation

Actors (N = 15; age range 20–62; 11 female) were recruited via opportunity sampling from UK drama schools. Approximately one week before filming, actors received a randomly ordered list of the 21 scripts, task instructions, and setup information. Actors were informed of the target emotion associated with each scenario but were not given intensity instructions or guidance on how to portray the emotion (e.g., no action-unit constraints). Actors were instructed to wear a black T-shirt, minimize facial makeup, look directly at the camera, and avoid turning away. They were asked to begin each portrayal with a neutral facial expression and develop the expression into the intended emotion over time.

### Recording setup and self-paced recording procedure

Recordings were conducted with actors seated on a stool against a black background. Video was captured using a Panasonic HDC-TM900 camera at 24 fps, mounted on a tripod at eye level approximately 70 cm from the actor. A laptop positioned approximately 60 cm in front of the actor displayed scripts and task prompts and was controlled via a customized keyboard used to advance trials and start/stop recordings.

Each trial proceeded as follows. A red fixation point and the word “neutral” prompted the actor to relax and maintain a neutral expression for 10 s. Next, one of the 21 scripts appeared, with the intended emotion category shown in parentheses. Actors read the script and prepared for as long as needed. When ready, the actor pressed a key that re-displayed “neutral” to enforce a neutral starting pose and simultaneously triggered video recording, marking the start of the preparation period. After 8 s, a 0.5 s medium-pitch beep signaled the actor to begin developing the expression into the intended emotion, marking the onset of enactment. Recording was terminated by the actor using the keyboard once they judged the portrayal to be clear and sufficiently long, marking the end of enactment. After each clip, actors completed brief self-report questions about their performance (including self-believability and emotional strength). For physiological analyses, the preparation epoch was defined as the interval from recording onset to beep onset, and the enactment epoch as the interval from beep onset to clip termination.

### Physiological recording during portrayals

To index cardiac dynamics during performance, we recorded electrocardiogram (ECG) continuously using a BioSemi system (BioSemi B.V., Amsterdam, The Netherlands) at a sampling rate of 1000 Hz. The electrodes were placed over the right clavicle and left iliac crest according to Einthoven’s triangular arrangement. These electrodes were positioned so they were not visible in the video frame. R-peaks were detected using MATLAB R2025a (MathWorks, Natick, Ma, USA) using the Signal Processing Toolbox (*findpeaks* function). Inter-beat intervals (IBIs) were computed from successive R–R intervals. These implementations were visually inspected. Artefact-contaminated segments were excluded on an actor-wise basis by removing IBIs outside ±1.5x the actor’s median IBI^47^. From the cleaned IBIs, we derived two summary measures per clip and epoch: mean heart rate (beats per minute) and heart-rate variability (HRV) quantified as RMSSD of successive IBIs. HR and RMSSD were computed separately for the preparation and enactment epochs. RMSSD is widely used in short recordings and can provide informative estimates over brief segments, although longer segments or repeated estimates can improve precision^48,49^.

### Video clip editing

Clips were edited using Shotcut video editor (Meltytech, LLC). Each clip was cut from the visual onset of the beep waveform (spectrogram onset) to approximately 0.5–1 s before the actor finished the portrayal; audio was muted. Clip length was allowed to vary to preserve naturalistic timing while remaining suitable for the study. Final clips were 6–13 s (mean 11 s) at 24 fps, with a 0.5 s black fade-in and fade-out to minimize orienting responses.

### Online validation study design

The full stimulus set comprised 311 clips, including 4 practice clips and 307 target clips. Of the 307 validation clips, 304 were included in the present analyses; three clips were excluded because one or more required measures (e.g., movement magnitude or clip-level ratings/covariates) were missing after preprocessing. To reduce participant fatigue, validation clips were divided into three fixed batches (A, B, C). Each batch contained 92 batch-specific clips plus 31 clips common to all batches (about 10% of the validation set) to enable cross-batch reliability assessment. Participants completed one survey corresponding to one batch and one rating dimension, yielding 12 surveys in total (3 batches × 4 dimensions). The four dimensions were emotion recognition (forced-choice identification of the portrayed emotion), valence, arousal, and believability, with scale anchors referring to whether the portrayal appeared real or natural. Recognition was included as a validation check to assess whether each clip was categorized as the intended emotion at the group level (alignment between the actor’s target emotion and observers’ categorization).

### Online validation participants and procedure

Participants were recruited in the UK via Prolific® and completed surveys implemented in Qualtrics®. A total of 404 individuals participated. After applying pre-specified quality criteria, we retained 371 participants who reported no problems viewing the clips and passed the engagement checks described below. Given the length of the whole stimulus set, participants were randomly assigned to one rating dimension; final retained sample sizes were: believability N = 93, valence N = 95, arousal N = 91, and recognition N = 92. Participants were instructed to complete the survey on a computer (not a mobile phone). Each batch-specific clip received at least 30 ratings per dimension; common clips accumulated ratings across batches. This target was set a priori to stabilize clip-level mean estimates and to provide adequate precision for stimulus-level analyses and mixed-effects models.

To assess engagement, each survey included two catch trials consisting of short, animated cartoon clips (about 5 s) depicting emotions at extreme ends of the relevant rating continuum. Catch-trial clips were selected from a pool of eight candidates based on ratings from an additional Qualtrics® sample, choosing the two clips with the highest Cronbach’s alpha per factor.

### Rating measures

Participants provided ratings for one of four dimensions. For recognition, participants selected which emotion was expressed from eight options (happiness, sadness, anger, surprise, fear, disgust, neutral, other). Clip recognizability was quantified as the proportion of observers selecting the actor’s intended emotion category (uncorrected); chance-corrected variants yielded the same qualitative conclusion. Valence and arousal were rated using continuous sliders from 0 to 100 (valence: very negative to very positive; arousal: very relaxed or calm to very excited or stimulated). Believability was rated using a continuous slider from 0 to 100 anchored as “not at all a real or natural emotion” to “very much a real or natural emotion.”

### Facial movement measures

To calculate a measure of facial movement, changes in the positions of facial landmarks over the videos were calculated. The OpenFace tool was used for facial landmark detection and alignment^50^. For each frame of the 304 video clips OpenFace returned the 2-D coordinates of 68 facial landmarks, the outer contour of the face, the eyebrows, the nose, the eyes, the mouth and the inner lips. For each marker, the position was averaged (using a sliding window) over 4 consecutive frames. The difference in position for each marker was calculated for consecutive frames and the magnitude of any change in position between frames was calculated. Frames in which there was a change between frames greater than 50 were excluded. This threshold was selected to identify rare frames in which OpenFace failed to correctly place the landmarks on the face. The remaining data were averaged across markers and across all frames to produce one number per video that represented the average movement magnitude across all markers. This value was used for all subsequent analysis and captured both head movements and changes in facial expressions.

### Quantification and Statistical Analysis Overview, outcomes, and data structure

Believability ratings (0–100) were the primary outcome. Valence and arousal ratings (0–100) indexed affective meaning at the clip level. Emotion recognition (forced choice) served as a validation check of alignment between intended emotion category and observers’ categorization. Analyses were conducted at two levels: stimulus-level analyses used clip-level means, and trial-level analyses used individual ratings while accounting for clustering by stimulus and observer (11,323 believability ratings from 93 observers across 304 clips). Trials with missing ratings were removed. Analyses were implemented in R (R Core Team. v4.5.1; packages included *lme4 v1*.*1-38, lmerTest v3*.*2-0, dplyr v1*.*2*.*0, tidyr v1*.*3*.*2, readxl v1*.*4*.*5*).

### Association between movement magnitude and believability

At the stimulus level, we quantified the association between movement magnitude and mean believability using Pearson correlation and linear regression. Movement magnitude, clip-level mean valence, and clip-level mean arousal were each z-scored across clips prior to entering the models, so all regression coefficients reflect a one standard deviation change. Intended emotion was included as a categorical covariate to assess whether movement effects persisted beyond emotion categories. At the trial level (11,323 ratings; 93 observers; 304 clips), believability ratings were analyzed using linear mixed-effects models (REML = FALSE) with random intercepts for observer and stimulus. Fixed effects included movement magnitude, intended emotion, and clip-level mean valence and arousal. Clip-level mean valence and arousal were derived from independent observer samples who rated those dimensions separately and did not contribute believability ratings. Nested model comparisons were evaluated using likelihood-ratio tests. To assess robustness, we refit the primary trial-level model with batch membership as an additional fixed effect, with stimulus-equalizing weights (1/n_ratings per stimulus), and with participant-specific slopes for movement magnitude; the movement effect remained negative and reliable across all specifications.

### Non-linearity and valence modulation

To test departures from linearity, we compared linear and quadratic specifications at the stimulus level and fitted spline-based trial-level models (natural cubic splines with 3 degrees of freedom) to capture non-linear movement effects. Valence modulation was tested by including a movement spline × valence interaction. Modulation by emotion category was examined as an exploratory nested comparison.

### Cardiac measures and believability

Cardiac physiology models examined two ECG-derived measures: mean heart rate (beats per minute) and heart-rate variability (HRV, quantified as RMSSD), computed separately for the preparation and enactment epochs of each clip. For heart rate, enactment values were expressed relative to each actor’s own mean enactment heart rate across all their clips (actor-mean correction) to remove between-actor baseline differences before testing for differences across intended emotion categories. For HRV, analyses were restricted to the 285 clips with complete data for both epochs. RMSSD was z-scored across clips before entering the models. Valence-dependent associations between HRV and believability were tested via HRV × valence interaction terms, and conditional slopes were estimated at one SD above and below the mean valence (valence_z = ±1) within the same model, without splitting the dataset. To verify that any HRV–believability association was not simply a downstream consequence of movement, we additionally fitted models that included a movement × valence interaction alongside the HRV × valence term. As a further robustness check, HRV models were refit with an additional random intercept for actor identity to account for potential clustering of clips within actors.

### Actor self-reports and mediation

Actor self-believability and emotional strength were related to observer believability at the clip level (N = 304). Mediation analyses tested whether any association between actor-rated emotional strength and observer believability operated indirectly through movement magnitude. The indirect effect was estimated as the product of two paths: path a, the association between emotional strength and movement magnitude (estimated via stimulus-level OLS regression controlling for clip-level valence, arousal, and intended emotion), and path b, the association between movement magnitude and believability controlling for emotional strength (estimated via trial-level LMM with random intercepts for observer and stimulus). The total effect of emotional strength on believability (c path) was tested via likelihood-ratio test prior to examining the indirect pathway. Cluster bootstrap resampling over stimuli was used to derive 95% confidence intervals for the indirect effect (B = 5000 iterations), resampling intact stimulus clusters together with all associated observer ratings to preserve the nested data structure.

## SUPPLEMENTAL INFORMATION

Document S1. Figures S1-3.

