## Supplementary material for "Actors’ Facial Movement Magnitude and Cardiac Dynamics Predict Observers’ Emotion Believability Ratings": Document S1

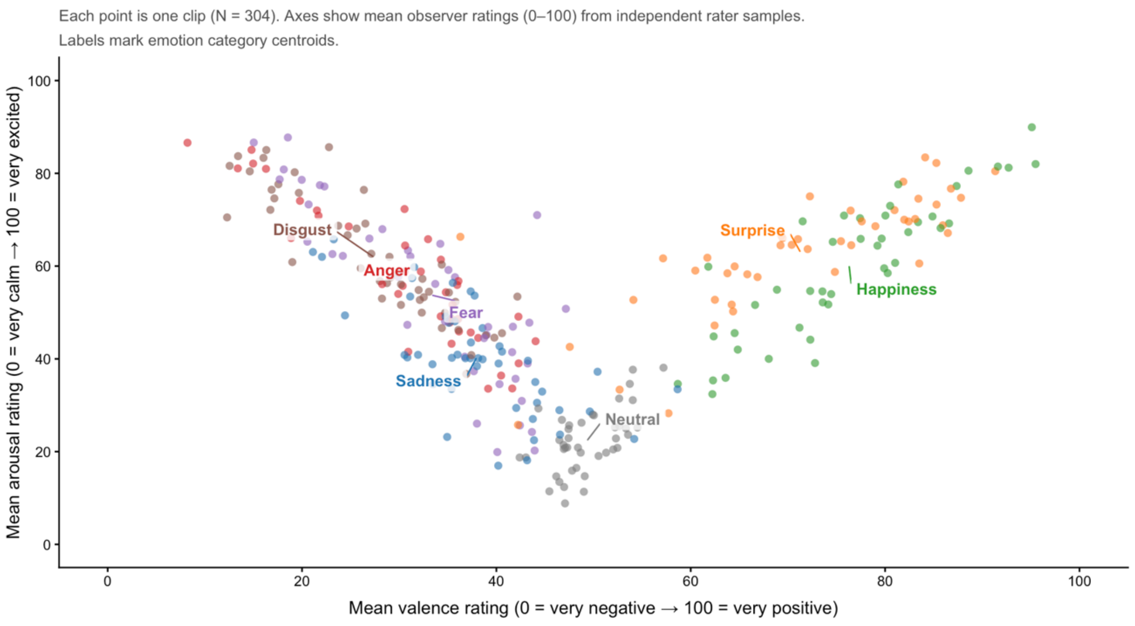


**Figure S1. Stimulus affective space by intended emotion.** Each point represents one analyzed clip (N = 304), plotted according to independent observer ratings of mean valence and mean arousal. Colors indicate intended emotion categories, and labels mark category centroids. The plot shows that intended emotions occupy distinguishable but partially overlapping regions of affective space.

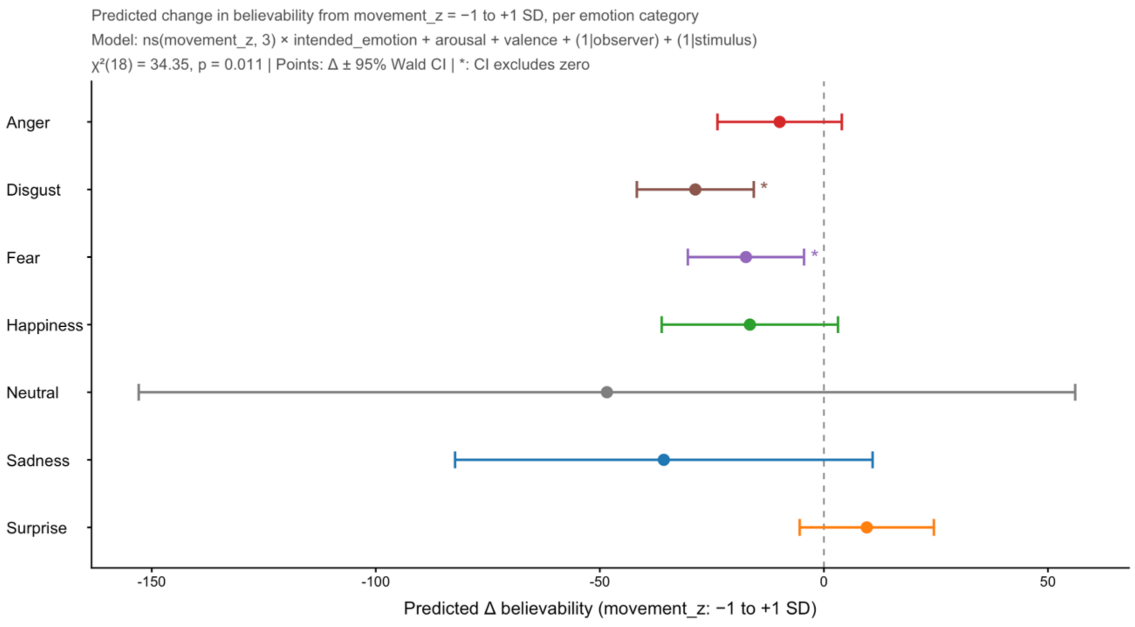


**Figure S2. Emotion-category modulation of the movement-believability slope.** Predicted change in believability from movement_z = −1 to +1 SD is shown separately for each intended emotion category. Points indicate estimated changes and error bars indicate 95% Wald confidence intervals from the trial-level spline mixed-effects model including movement × intended emotion, arousal, valence, and random intercepts for observer and stimulus. The model comparison indicated additional emotion-category modulation of the movement-believability mapping, $\chi^2(18) = 34.355$, $p = 0.011$.

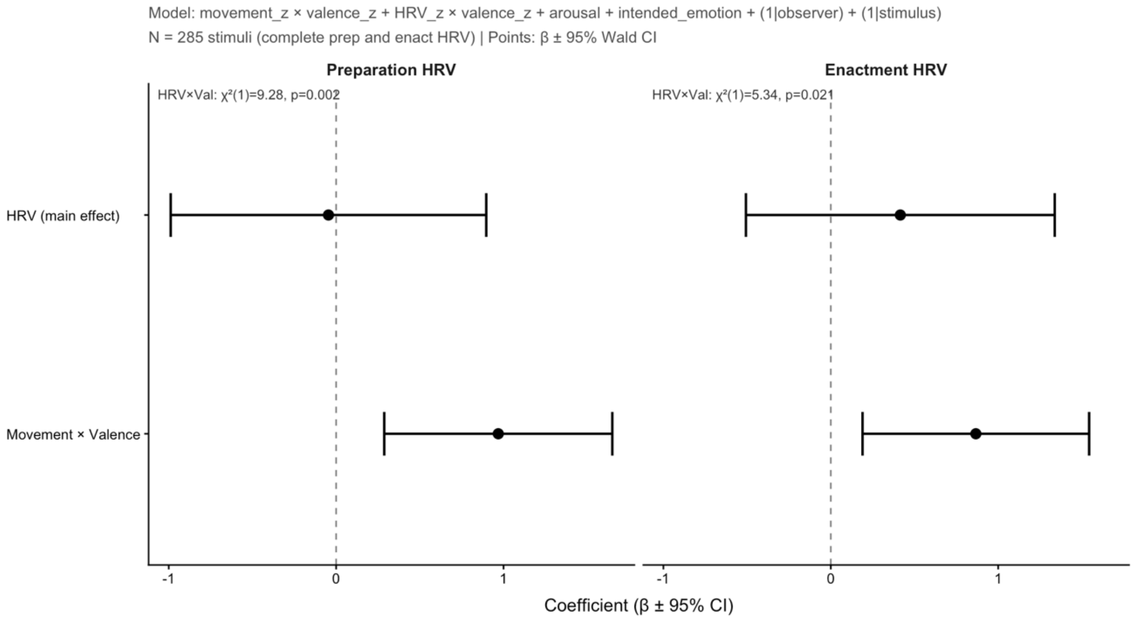


**Figure S3. HRV × valence effects after controlling for movement × valence.** Coefficient estimates from strict HRV models testing whether HRV × valence effects remained after including movement × valence terms. Points indicate fixed-effect estimates and error bars indicate 95% Wald confidence intervals. HRV × valence remained reliable for both preparation HRV, $\chi^2(1) = 9.28$, $p = 0.002$, and enactment HRV, $\chi^2(1) = 5.34$, $p = 0.021$, indicating that the HRV-believability association was not reducible to the valence-dependent movement effect.
